# Towards extracting supporting information about predicted protein-protein interactions

**DOI:** 10.1101/031591

**Authors:** Adam Roth, Sandeep Subramanian, Madhavi K. Ganapathiraju

## Abstract

One of the goals of relation extraction is to identify protein-protein interactions (PPIs) in biomedical literature. Current systems are capturing binary relations and also the direction and type of an interaction. Besides assisting in the curation PPIs into databases, there has been little real-world application of these algorithms. We describe UPSITE, a text mining tool for extracting evidence in support of a hypothesized interaction. Given a predicted PPI, UPSITE uses a binary relation detector to check whether a PPI is found in abstracts in PubMed. If it is not found, UPSITE retrieves documents relevant to each of the two proteins separately, and extracts contextual information about biological events surrounding each protein, and calculates semantic similarity of the two proteins to provide evidential support for the predicted PPI. In evaluations, relation extraction achieved an Fscore of 0.88 on the HPRD50 corpus, and semantic similarity measured with angular distance was found to be statistically significant. With the development of PPI prediction algorithms, the burden of interpreting the validity and relevance of novel PPIs is on biologists. We suggest that presenting annotations of the two proteins in a PPI side-by-side and a score that quantifies their similarity lessens this burden to some extent.

## 1. Introduction

Protein-protein interactions (PPIs) play an important role in the understanding of human biology. By analyzing PPIs and their interaction networks, researchers can unravel the molecular mechanisms of disease progression [1]. Discovery and exploration of such interactions is fundamental to biological, pharmaceutical, and medical research [2][3]. The BioGRID interaction database currently contains 19,595 unique proteins and 185,112 non-redundant human PPIs [4]. Current studies have estimated the total size of the human interactome to be ~650,000 interactions, leaving an estimated 500,000 undiscovered interaction pairs [5].

**Wet-lab discovery of all individual interactions is unrealistic using current detection methods**. High-throughput techniques such as yeast two-hybrid (Y2H) are plagued by high false-negative and high false-positive rates of up to 70% [6][7]. High confidence methods such as co-immunoprecipitation have low-throughput, and are of significantly high cost in terms of time, effort, and money [8][9]. Computational discovery of the interactome has thus become a priority in bioinformatics as a method to guide biology research. Supervised learning for PPI prediction utilizes information on known PPIs to produce manageable subsets of plausible interactions. A number of supervised machine learning algorithms have been applied to PPI prediction including support vector machines[10], decision tree [11], Bayes classifier [12], kernel-based [13], and random forest based methods [14]. These approaches analyze patterns in known biological information to infer high-confidence interactions. Prediction methods are commonly placed into six categories based on input data including protein sequence, protein structure, genomic context, homology, experimental profiles, and literature-derived associations [15]. The computational and statistical mechanisms of supervised learning algorithms are well understood by machine learning specialists, but wet-lab researchers often have difficulty accepting output as valid indication that an interaction is likely to occur. Justification of predicted PPIs is thus paradoxically tasked to the wet-lab researcher who must locate relevant information in the scientific literature.

**Our collective knowledge of protein functions and pathways is scattered somewhere among 23 million articles in PubMed** [16]. The number of articles continues to grow at the rate of 2 documents per minute [17], and it has become increasingly difficult for scientists to locate and distill relevant information [18]. Information collation through manual literature search has thus become a bottleneck in the process of scientific discovery [19]. The field of text mining has had a recent focus on the analysis of biomedical literature with the intent to enable scientists to harness the information available within PubMed. Many different approaches have been developed to extract information regarding PPIs. Such approaches include machine learning systems as well as empirical rule-based information extraction systems [20]. Complex events are typed n-ary associations of entities or other events that are characterized by an event trigger (typically a verb indicating an action) and one or more event participants [21]. Event participants can be either named-entities or other events and play specific roles in the event such as *Theme* or *Cause.* Specifically, complex event extraction extends binary relation extraction by identification of additional information about a given relation including direction, interaction type, binding site, and argument nesting [22]. Automated text mining efforts have been evaluated at the PubMed scale. The Turku Event Extraction algorithm (TEEs) developed by Bjourne et al. achieved state-of-the-art performance of 50.06% recall, 59.48% precision, and 54.37% F-score on the BioNLP ‘09 dataset [23][24]. It was approximately generalizable for large-scale application. When applied at the PubMed scale, TEEs extracted 21.3 million detailed bio-molecular events including protein-protein interactions [25].

**Various approaches have been studied to apply biomedical text-mining to PPI prediction**. In general, these methods are similar to non-text mining PPI prediction approaches in that they use data on known interactions to generate interaction hypotheses. Binary relation extraction has been utilized to generate interaction networks and uncover interactions hidden in free-text. Such methods are promising because they expand coverage of the known proteome [15] and can significantly accelerate PPI database curation efforts [26][27]. The topology of automated interaction networks generated through text-mining can be exploited for use in interaction prediction [28]. Network models based on the co-occurrence of proteins within sentences, abstracts, or articles have been found to reflect functionally relevant relationships [29]. Analysis of co-occurrence networks has shown that proteins and genes co-mentioned in the same article can reasonably be assumed to be related in some way [30][31]. Co-occurrence analysis has also been applied to connect proteins to diseases, biological processes, phenotypes, chemicals and key words found in articles [32][30]. Hidden and unknown biological associations can be predicted with high confidence using co-occurrence based methods. The degree of co-occurrence can be quantified to eliminate statistically weak associations and increase prediction accuracy [31][15].

**Hidden relationships between two proteins can be confirmed by mining the literature for shared concepts**. If we assume that A and C are both related to B, A and C may also share a direct relationship, and literature mining may reveal the shared concepts between A and C [33]. This method of network analysis has been applied to PPI prediction in various ways. Even if two proteins do not have an explicitly defined relationship in the literature, a functional relationship can be extrapolated by connecting both proteins to an overlapping set of intermediate concepts (B_1_-B_n_). Intermediate concepts are often designated to be the neighbors of A and C in an interaction network graph, but studies have shown success in linking proteins through intermediate concepts such as disease [34], keywords [35], biological processes [13], orthology [36], and user-defined biological terms [34]. Novel relationships have been extracted from the literature in this way and confirmed to have biological relevance [32][37]. It has been shown that concepts shared in A-B and B-C relationships can be located in the literature an average of 6.5 years in advance of the first explicit appearance of an established A-C relationship [33]. This time lag serves as an indication of the potential to which text-mining aided PPI prediction can accelerate wet-lab discoveries.

**Verbs are important to mine interaction networks because they are a syntactical requirement for expressing relations**. PPI networks can be organized using triplet representation where the proteins are nodes and verbs are edges between nodes. Cohen et al. showed that alterations in the argument structure of verbs and their nominalizations are both common and exceptionally diverse throughout subdomains of biomedicine [38]. Hierarchical clustering of verb sub-categorization throughout the scientific literature has been shown to form stable clusters that represent various biomedical sub-domains [39]. Small pockets of specialized behavior in verb subcategorization have been noted in addition to many cases of specific usage within a single subdomain [40].

**We present UPSITE, a text mining algorithm to extract information that supports or signifies a computationally predicted interaction**. There is little research on the application of text mining to *support a hypothesized PPI* as opposed to extracting factual information that is directly reported in literature. Given a protein pair (A and C) that is hypothesized to be an interacting pair, UPSITE examines the scientific literature and extracts sentence level information which assists in human interpretation of the interaction plausibility. Complex event extraction is used to identify the mutual concepts of A-B and C-B where mutual concepts are event triggers extracted from the triplet representation of mined interactions. Event triggers are used as mutual concepts due to their high level of importance to interaction identification as well as their ability to accurately segregate biomedical subdomains (Figure 1). UPSITE can be used to identify direct as well as indirect information that supports known or predicted interactions, respectively.

**Figure 1.**
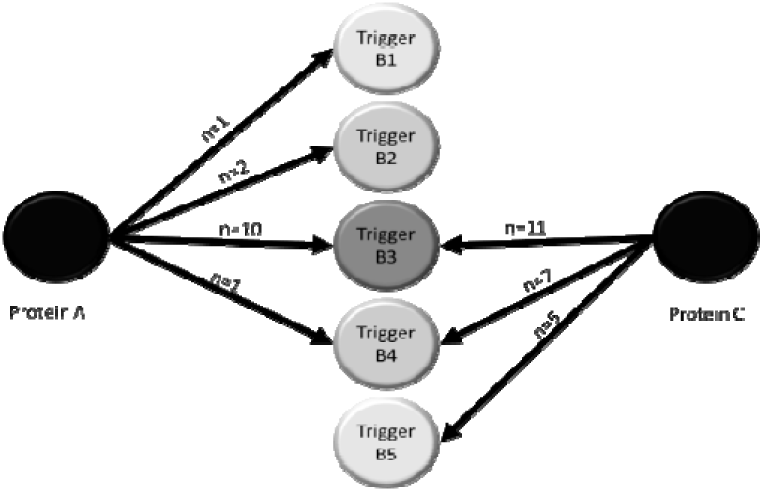
Measuring semantic similarity between proteins.

## 2. Methods

We have developed UPSITE, a text mining tool for validating interaction hypotheses. The UPSITE pipeline consists of three main steps: 1. relation extraction; 2. complex event extraction; and 3. semantic similarity. In this section, we provide a detailed description of the UPSITE workflow. Input to UPSITE is a pair of proteins (also called a query pair) which have been predicted to interact. UPSITE is designed to enable the exploration of the relationship between the proteins in the query pair.

### Information Retrieval

Query expansion refers to the process of reformulating a seed query to improve information retrieval (IR). Queries and synonyms are normalized by removing non-alphanumeric characters. Query expansion is then performed using a synonym database constructed from the gene symbols contained within 6 genomic and proteomic databases by Chen and Sharp [41]. The expanded query is used to construct an Entrez URL-query according to the NCBI Entrez Programming Utilities documentation [42]. PubMed abstracts are obtained in XML format through Entrez *esearch* and *efetch.*

### Relation Extraction

Relation extraction refers to the natural language processing (NLP) techniques that identify a semantic relationship between terms. UPSITE is designed to identify both explicit and implicit PPIs. A sentence contains a relationship of interest if it describes or infers any influence or biologically relevant correlation between queried proteins. This definition is relaxed from that of an explicit interaction in order to increase recall of extracted sentences. Unlike conventional binary relation extraction systems which focus on extracting explicit interactions, UPSITE extracts any sentences that are potentially useful in verifying an interaction. Predicted interactions are not likely to be described in the current scientific literature. We thus focus on identification of sentences that support interaction plausibility.

We choose single sentences as our unit of analysis [43]. Only sentences containing co-occurrence of the query proteins are further processed. Queries and their synonyms are normalized by removing all non-alphanumeric characters. Filtering is performed by running a case-insensitive search for the normalized queries and their synonyms. Sentences that do not contain both proteins are disregarded. Prior to the relation detection phase, parentheses and parenthetical contents that do not contain proteins are discarded. Query proteins are blinded and replaced with ‘Protein1’ and ‘Protein2’.

Our method of relation detection involves two scoring modules which assign scores according to empirically determined rules. We will refer to the scoring systems as ‘scoring system 1’ and ‘scoring system 2’, reflecting their order in the program flow. The remaining sentences are part-of-speech (POS) tagged with the NLTK POS tagger [44]. Scoring system 1 extracts sentence features according to a rule-based, template matching system. Features were selected by conducting a review of the current text mining literature. The score assigned to each attribute was determined by its frequency of occurrence in a dataset of 200 randomly chosen sentences from the BioCreative II PPI dataset [45]. Lists of trigger words were generated by curation of verbs indicative of stimulation, conclusion, or inhibition. A sample of individual attributes and their respective scores can be found in Table 1.

**Table 1.** Example scoring attributes of scoring module 1

| Score | Attribute |
| --- | --- |
| +3 | Exact query term in sentence |
| +5 | First word is query term |
| +20 | First word is query term and indicator of stimulation in sentence |
| +5 | Second word is a verb |
| +5 | Method of testing for interaction is mentioned |
| +5 | Indicators of Stimulation |
| +9 | Indicators of Conclusion |
| -3 | Indicators of Failure |
| -2 | Long sentences (>30 words) |
| +10 | Protein complex is described (Protein1-Protein2) |
| +5 | Stimulation word in sentence |

Positively scored sentences from the scoring system 1 are then passed to scoring system 2 which makes further use of NLP to decide the final sentence rankings. While scores from module 1 indicate the presence of a biological relation (syntax), scoring system 2 deciphers whether or not the query terms are involved in the relation (semantics). A parse tree is an ordered, rooted tree that represents the syntactic structure of a sentence [46]. A statistical parser trained on a Penn Treebank probabilistic context-free grammar was used to generate parse trees using the Cocke–Younger–Kasami (CYK) parsing algorithm [47]. The parse tree is traversed using preorder traversal (depth-first search) to create an ordered list of simple declarative clauses, verb phrases, noun phrases, query terms, and their location indices. Sentences are then assigned scores based on relative location of query terms and their Verb Phrase Verbs. A flowchart of UPSITE relation extraction can be found in Figure 2.

**Figure 2.**
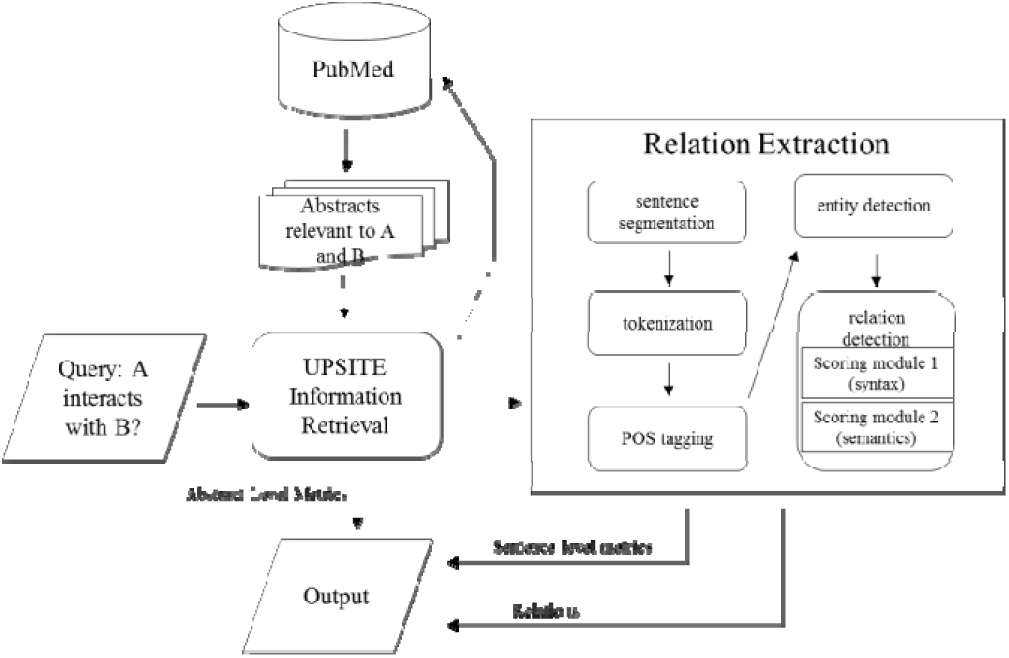
Relation extraction workflow

The tag for a noun phrase in a parse tree is NP and the tag for the noun of a noun phrase is NN (Noun, singular or mass), NNS (Noun, plural), NNP (Proper noun, singular), or NNPS (Proper noun, plural). Figure 3 highlights the importance of the noun phrase in detecting inferred interactions. In Figure 3, the noun phrase “the expression of LATS2 and MDM2, hTERT and MDM2” contains both of the proteins of interest (MDM2 and hTERT) within a single complex noun phrase (indicated by NX). Preorder traversal of the parse tree is able to locate this close relationship between these two proteins. Similarly, an example verb phrase describing an interaction between thrombin and cd69 is, “Thrombin induces tcr cross-linking for cd69 expression and interleukin 2 production”. Sentences containing relevant noun phrases and verb phrases are weighted +20. UPSITE outputs the top ten highest scoring sentences.

**Figure 3.**
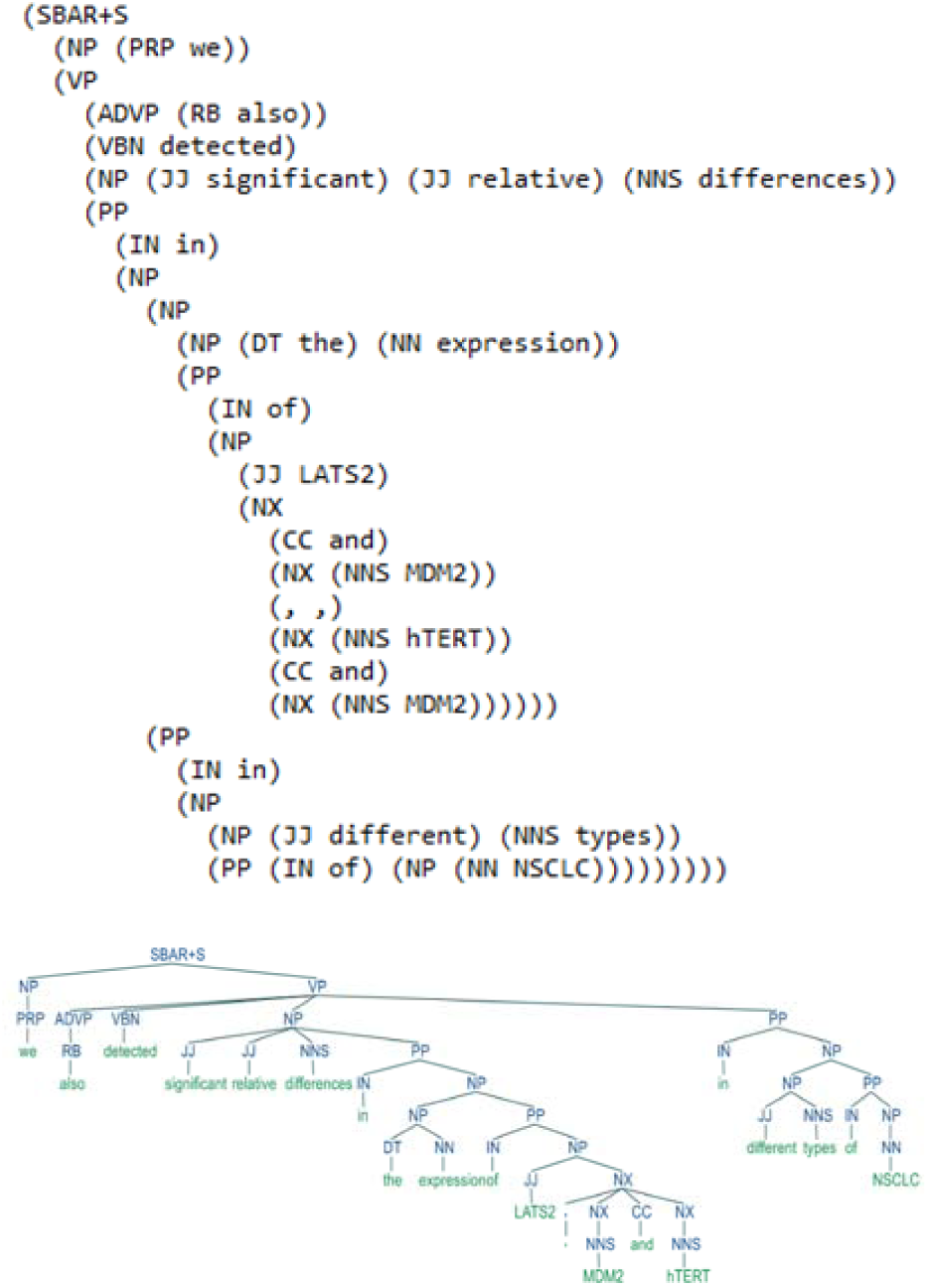
Example of parse tree

### Complex Event Extraction

Given a pair of proteins (protein-A and protein-B), if they are not known to interact, an explicit statement of interaction is unlikely to occur in published literature. The relation detection module of UPSITE attempts to extract sentences implying or speculating interaction, but this method is limited to the sentence as a unit of analysis. We hypothesize that this barrier can be broken through directed collation of relevant biological knowledge across PubMed. To test this hypothesis, we have developed a method to generate support for undocumented PPIs by analyzing textual descriptions of known interactions relevant to the query proteins. The queried pair is first split into its constituent proteins. Each query protein is separately processed through the event extraction module. UPSITE’s information retrieval module was modified to retrieve abstracts relevant to a single protein and its synonyms. Retrieved abstracts are then processed by the TEES complex event extraction algorithm. The TEES code is open source and freely available. We modified the TEES code to enable bulk processing of retrieved abstracts. Processing is completed using the GE11 training model which was developed for the BioNLP’11 shared task [23]. The output of TEES is a set of interaction XML files containing complex events and their respective trigger words. UPSITE parses the interaction XML files and generates a list of trigger words. Word lists include triggers for all interactions in the set of retrieved abstracts relevant to a given query protein. Following generation of trigger word lists, a distance metric is computed to measure their similarity. A flowchart of UPSITE semantic similarity can be found in Figure 4.

**Figure 4.**
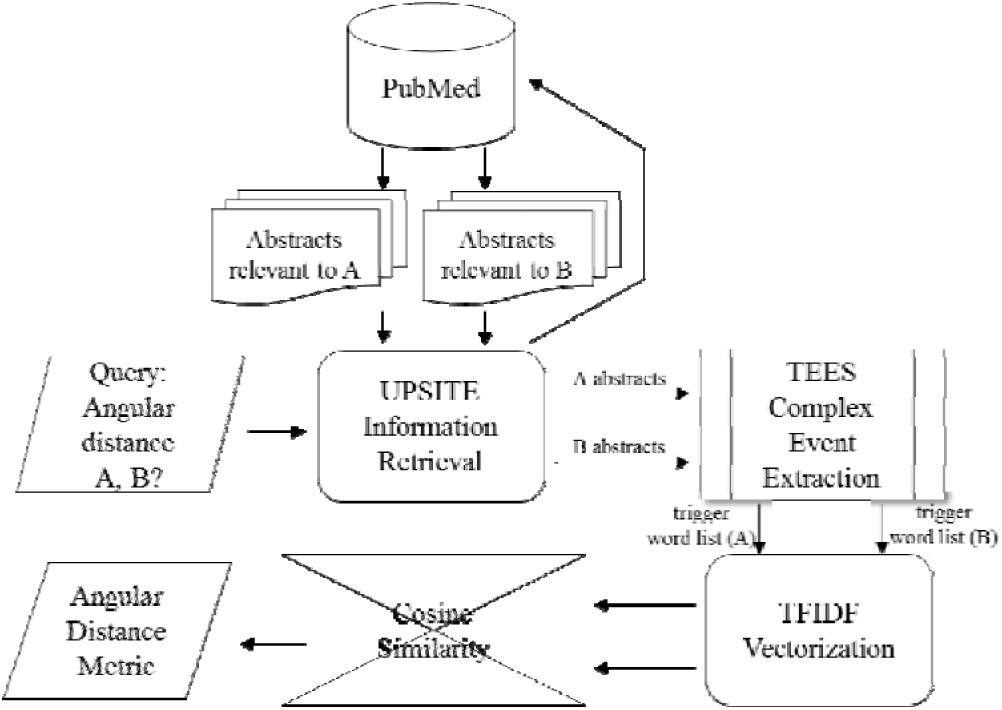
Semantic similarity workflow

### Semantic similarity

Semantic similarity of trigger word lists is measured by cosine similarity. To obtain this measurement, a document-term matrix is formed using term frequency-inverse document frequency (tf-idf) vectorization as a function of cell counts. Cosine similarity is obtained using the linear kernel which is equivalent to the cosine similarity measure ——— [48]. Cosine similarity was chosen as our distance function based off of a study by Xiang et. al comparing various methods of comparing vectors including dices coefficient, horn coefficient, euclidean distance, and manhattan distance and found cosine similarity to be the most accurate measure [49]. The cosine measurement is then converted to angular distance for ease of interpretability by first converting to radians, then converting to degrees.

## 3. Results

### Textual relationship validation

The HPRD50 corpus was used to benchmark the performance of UPSITE relation extraction. Although UPSITE differs from standard PPI binary relation detection algorithms in that it has been optimized to detect implied rather than plainly expressed interactions, we felt that a comparison to current methods would still be a useful indication of system performance. The HPRD50 dataset was developed by Fundel et al. as a test set for the RelEx relation extraction system [50]. It contains sentences from 50 abstracts referenced by the Human Protein Reference Database (HPRD)[51] and includes 145 annotated interactions [52]. We chose the HPRD50 dataset because it defines explicit interactions (e.g. direct physical interactions) in addition to implicit interactions (e.g. regulatory relations and modifications) [50]. To optimize performance against this dataset, UPSITE’s rule-based scoring system was used to generate feature vectors for a support vector machine (SVM). Individual features were weighted according to the respective scores assigned in UPSITE’s scoring module. The entire algorithm was implemented using Python and a python wrapper to TEES. SVM was implemented using the Python library Scikit-learn [53]. UPSITE achieved 0.88 F-score, 0.94 precision, and 0.83 recall. The performance of UPSITE was compared to a similar algorithm (RelEx) as well as a baseline standard of sentence level co-occurrence. UPSITE demonstrated a 10% increase in F-score as compared to RelEx and 39.6% increase as compared to baseline co-occurrence. Performance comparison can be seen in Table 2. UPSITE was run using the Ubuntu 14.02 operating system with a 2.5 Ghz Intel Core i7–4710HQ and 16GB RAM. Running time averaged to 3.1 hours per protein pair. UPSITE was then tested against the BIONLP shared task 09’ corpus. This test set included protein interactions which were manually curated from the abstracts of PubMed articles and is available online. We manually analyzed results on 200 interactions and achieved .803 recall, .901 precision, and .849 F-score.

**Table 2.** – Performance of Binary Relation Detection methods on HPRD50 corpus

|  | Precision | Recall | F-score |
| --- | --- | --- | --- |
| UPSITE | 0.94 | .0.83 | 0.88 |
| RELEX algorithm | 0.80 | 0.80 | 0.80 |
| Co-occurrence | 0.46 | 1 | 0.63 |

### Cosine similarity and complex event extraction

Complex event extraction was performed by the TEES algorithm. The extraction performance of TEES has been extensively evaluated on the BioNLP Shared Tasks and it was the winning system of ST’09 achieving 58.28% precision, 46.73% recall, and 51.96% F-score. TEES has since been updated to TEES 2.2 and shown to achieve state-of-the-art results in ST’11 with F-score 54.37%, precision 59.48%, and recall 50.06% [25]. Although not used in this experiment, further modification of the TEES system using feature selection has pushed its performance to an F-score of 57.24% [54].

50 protein pairs known to interact (known) and 50 randomly assigned protein pairs (random) were used to analyze performance and usefulness of the angular distance metric for trigger word lists. Average angular distance of known and random pairs was measured as a function of the number of papers parsed. Figure 5 shows the results of this analysis. Both known and unknown pairs were shown to closely follow a logarithmic regression with respective coefficients of determination of 0.9865 and 0.9633. A two-sample Analysis of Variance (ANOVA) revealed an F-statistic of 1.284 and p-value of 0.34. Thus, the observed variances are not statistically different. A two-tailed, two-sample t-test assuming unequal variances revealed a p-value of 0.002024 and T-statistic of −3.4677. Because the observed means were not significantly different, we conclude that the angular distance metric is statistically valid at *α* = 0.005. ANOVA and T-test can be found in tables 3 and 4. In summary, angular distance metrics were calculated for 50 pairs of proteins known to interact (known dataset) and 50 randomly paired protein (random dataset). We compared the distance scores using a two-tailed, two-sample t-test assuming unequal variances. The results of this test indicate that the scores from the known and random dataset are significantly different. The small p-value verifies that the method outlined in this study is useful in predicting whether two proteins are likely to interact.

**Figure 5.**
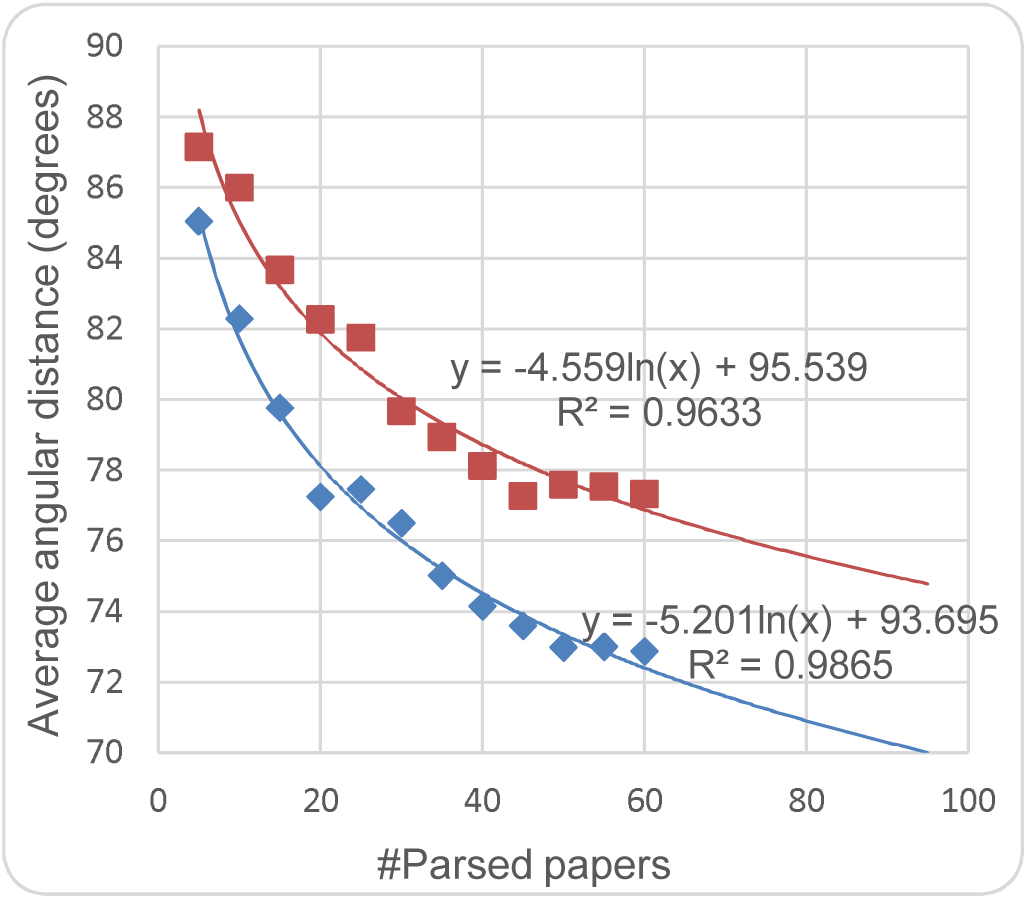
-Comparison of average angular distance

**Table 3.** -F-test Two-Sample for Variances

|  | Known Pairs | Random Pairs |
| --- | --- | --- |
| Mean | 77.6 | 84.74163 |
| Variance | 18.1 | 14.05495 |
| Observations | 12 | 12 |
| df | 11 | 11 |
| F | 1.2 |  |
| P(F<=f) one-tail | 0.34 |  |
| F Critical one-tail | 2.81 |  |

**Table 4.** -t-Test Two-Sample Assuming Unequal Variances

|  | Known Pairs | Random Pairs |
| --- | --- | --- |
| Mean | 77.6 | 84.74 |
| Variance | 18.05 | 14.05 |
| Observations | 12 | 12 |
| Hypothesized Mean Difference | 0 |  |
| df | 22 |  |
| t Stat | -4.363775 |  |
| P(T<=t) one-tail | 0.0001241 |  |
| t Critical one-tail | 1.7171444 |  |
| P(T<=t) two-tail | 0.0002481 |  |
| t Critical two-tail |  |  |

### Availability

We propose to make the supporting information extracted by UPSITE for each PPI on Wiki-Pi [57], a web-server that presents information about individual PPIs, in future. We have made the tool available as an AMI (Amazon Machine Instance) with all software dependencies, TEES models, corpora and NLP tools already setup. Detailed instructions on launching the AMI can be found on our GitHub repository^1^. The entire source-code has been made available on the same repository with an installation script that downloads and installs all of the necessary software dependenices. The tool can be run via the command line using command line arguments that include the pair of proteins to be analyzed, the number of papers to be parsed per protein etc. Results may be written to an output file that contains the similarity score between the protein pair and related terms extracted from literature.

## 4. Discussion

We have created UPSITE, a text mining algorithm for extracting supporting information about predicted proteinprotein interactions from scientific literature. Input to UPSITE is a pair of proteins or a list of protein pairs. UPSITE assists determination of interaction plausibility by extracting relevant textual data from the PubMed corpus. This algorithm was designed to address the gap between discovering a PPI computationally and choosing to validate it experimentally. While PPI prediction algorithms have the potential to accelerate wet-lab research, many biologists remain apprehensive about their use. This lack of confidence is not surprising given the cost of resources that are to be invested in studying a predicted PPI in the lab. UPSITE fills this information-void by providing textual evidence to support interaction hypotheses.

While previous studies have established the usefulness of semantic similarity in characterizing biological relationships, the input data for these studies have been manually curated from scientific literature or extracted from preexisting databases [55][56]. We have shown that useful input data for comparing semantic similarity of proteins can be automatically extracted from free text to accurately classify protein relationships. UPSITE can perform directed semantic similarity by selectively extracting trigger words for inclusion in the vector space model and calculating angular distance between query vectors.

The performance of relationship extraction and semantic similarity is restricted by the amount of descriptive information contained in the scientific literature. Yu et al found that the majority (83%) of currently known human PPIs have been cited only once [58]. Biomedical text mining is thus often faced with a large amount of data sparsity. This is especially an issue when dealing with predicted interactions because they are highly unlikely to have been previously verified in the literature. In small scale testing (100 protein pairs), we found single sentence relation extraction to provide useful validation evidence in 43% of test cases. However, cosine similarity was able to provide useful validation evidence in 95% of test cases. These data suggest that restriction of relation extraction to only co-occurrence sentences may constrain system performance.

In future work, we hope that the methods described in this paper can be expanded upon by further harnessing complex event extraction to improve directed semantic similarity complex event extraction to improve directed semantic similarity. Complex event extraction methods have been developed for the mining of various data types including interaction location, pathway, and relatedness. This information could be integrated into UPSITE to improve directed semantic similarity. Furthermore, future research should implement and compare performance of various similarity metrics including the Jaccard Index, Euclidean distance and latent semantic analysis.

In sum, UPSITE is able to shorten the gap between computer prediction and wet-lab verification of interacting protein pairs by providing textual validation of PPI hypotheses. PPIs are fundamentally important to the human biological system and their discovery can have effects throughout biological, medical, and pharmaceutical research. We hope that UPSITE will help to drive the adoption of PPI prediction algorithms by increasing wet-lab researchers’ confidence in machine learning predictions.

## 6. Acknowledgment

The authors thank Dr. Tanja Bekhuis and Katrina Romagnoli for their feedback during the writing of this paper. This work has been funded by the BRAINS grant R01MH094564 from the National Institute of Mental Health of the National Institutes of Health (NIMH/NIH), USA. All correspondence should be addressed to M. K. Ganapathiraju.

**Adam Roth** obtained his B.S. in Molecular Biology from the University of Pittsburgh, Department of Biological Sciences in 2012 and M.S. (expected) from the University of Pittsburgh, Department of Biomedical Informatics in 2015 from University of Pittsburgh, USA. He currently works for the U.S. Department of Veterans Affairs Informatics and Computing Infrastructure

**Sandeep Subramanian** obtained his B.S. in Computer Science and Engineering from the VIT University, India in 2014. He is currently a graduate student and a research assistant at the Language Technologies Institute at Carnegie Mellon University, Pittsburgh USA pursuing an M.S in Language Technologies.

**Madhavi K. Ganapathiraju** obtained B.Sc degree from Delhi University, India, M.Engg degree from Department of Electrical Communications Engineering, Indian Institute of Science, and Ph.D. in Language and Information Technologies from School of Computer Science, Carnegie Mellon University in 2007. She is currently Assistant Professor at Department of Biomedical Informatics with Secondary Appointment in Intelligent Systems Program, University of Pittsburgh. She holds affiliated faculty position at Language Technologies Institute, Carnegie Mellon University. She is *Member* of IEEE.

## Footnotes

1 https://github.com/MaximumEntropy/UPSITE.git

